# Machine Learning-based Biomarkers Identification and Validation from Toxicogenomics - Bridging to Regulatory Relevant Phenotypic Endpoints

**DOI:** 10.1101/2020.12.18.423486

**Authors:** Sheikh Mokhlesur Rahman, Jiaqi Lan, David Kaeli, Jennifer Dy, Akram Alshawabkeh, April Z. Gu

## Abstract

High-throughput in vitro assays and AOP-based approach is promising for the assessment of health and ecotoxicological risks from exposure to pollutants and their mixtures. However, one of the major challenges in realization and implementations of the Tox21 vision is the urgent need to establish quantitative link between *in-vitro* assay molecular endpoint and *in-vivo* phenotypic toxicity endpoint. Here, we demonstrated that, using time series toxicomics *in-vitro* assay along with machine learning-based feature selection (MRMR) and classification method (SVM), an “optimal” number of biomarkers with minimum redundancy can be identified for prediction of phenotypic endpoints with good accuracy. We included two case studies for *in-vivo* carcinogenicity and Ames genotoxicity prediction with 20 selected chemicals including model genotoxic chemicals and negative controls, respectively, using an *in-vitro* toxicogenomic assay that captures real-time proteomic response data of 38 GFP-fused proteins of *S. cerevisiae* strains covering biomarkers indicative of all known DNA damage and repair pathways in yeast. The results suggested that, employing the adverse outcome pathway (AOP) concept, molecular endpoints based on a relatively small number of properly selected biomarker-ensemble involved in the conserved DNA-damage and repair pathways among eukaryotes, were able to predict both *in-vivo* carcinogenicity in rats and Ames genotoxicity endpoints. The specific biomarkers identified are different for the two different phenotypic genotoxicity assays. The top-ranked five biomarkers for the *in-vivo* carcinogenicity prediction mainly focused on double strand break repair and DNA recombination, whereas the selected top-ranked biomarkers for Ames genotoxicity prediction are associated with base- and nucleotide-excision repair. Current toxicomics approach still mostly rely on large number of redundant markers without pre-selection or ranking, therefore, selection of relevant biomarkers with minimal redundancy would reduce the number of markers to be monitored and reduce the cost, time, and complexity of the toxicity screening and risk monitoring. The method developed in this study will help to fill in the knowledge gap in phenotypic anchoring and predictive toxicology, and contribute to the progress in the implementation of tox 21 vision for environmental and health applications.

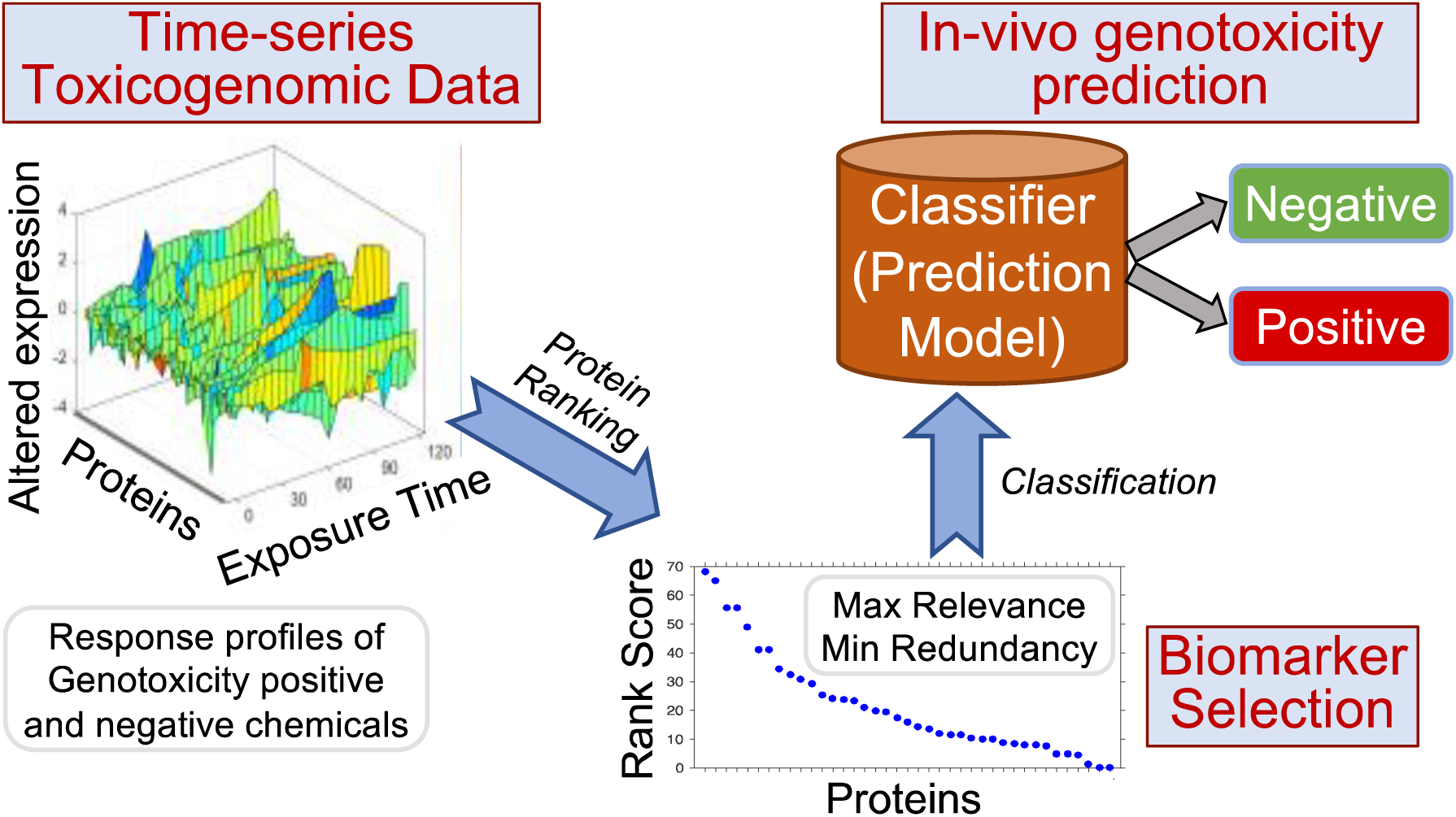
TOC Art

## INTRODUCTION

Genotoxicity is of great concern because of its link to mutagenicity, carcinogenicity as well as cancer, and there is urgent demand for genotoxicity screening and risk assessment for various environmental and health applications.^1-4^ In the absence of, or combined with, *in-vivo* carcinogenicity data, *in-vitro* or cell-based genotoxicity assays provide supporting data for cancer risk assessment.^5^ Recently, toxicogenomics has emerged to be a promising technology that reveals molecular-level activities, at the gene, protein, or metabolite level of organisms, in response to environmental contaminants and may represent the underlying cellular network mechanisms of toxicity responses.^2,6^ This also responds to the Tox21 vision that promotes a systematic transit from current *in-vivo* whole animal-based testing, to more *in-vitro* mechanistic pathway-based assays using high-throughput screening and tiered testing.^7-8^ However, one of the major challenges in realization and implementations of the Tox21 vision is the urgent need to establish quantitative link between *in-vitro* toxicogenomic assay molecular endpoint and *in-vivo* phenotypic regulatory relevant endpoints.

Establishing quantitative causal relationships between *in-vitro* assay endpoints to regulatory-relevant apical endpoints holds the key to the realization of predictive toxicology through practical and widespread implementation of *in-vitro* assay-based toxicity screening schemes and strategies for environmental and health applications.^9-16^ Adverse-outcome pathway (AOP) framework is the state-of-the-art approach to link mechanistic toxicity mechanisms with the phenotypic adverse outcome that would enable the assessment of health risk as well as ecotoxicological risks from exposure to pollutants and their mixtures.^17-21^ Coalesce of effective biomarkers and proper predicting framework would enable more cost-effective and wider implementation of toxicomics in monitoring of genotoxicity and predict adverse toxic responses.^22- 24^ Subsequently, proper selection and validation of predicative biomarkers plays a crucial role in our ability to link molecular-level effects recorded in *in-vitro* assays to the *in-vivo* regulatory relevant phenotypic endpoints, or system-level impacts in many fields such as, environmental toxicity, disease prediction and health risk identification.^25-28^

The rapid advancement in bioinformatics and machine learning methods enables more sophisticated biomarkers identification.^10,29-30^ Current biomarker identification from toxicomics data employs feature selection and classification methods. Two general approaches of feature selection include filter and wrapper methods. The filter methods often provide relatively simpler and faster alternatives to select the most important features and the features are selected or filtered based on their relevance to differentiate a target outcome from others.^31^ The wrapper methods combine feature selection along with the classification method, where the features are judged based on their ability to increase the accuracy of the classification models.^32-33^ However, the wrapper methods are often associated with extensive computational cost, and prone to possible overfitting when the sample size is relatively small.^34-35^ In addition, since the selected features of the filter methods are independent of the classification method, they often have higher relevance to the target outcome than those derived from a wrapper method.^31,34^ Filter based feature selection methods that have been applied to toxicomics data (e.g., gene and protein expression data) include mutual information, statistical tests (t-test, F-test, chi-square), information gain,^36^ gainratio,^30^ and ReliefF^36^ among others. Though most of these algorithms find the important biomarkers based on their relevance and correlation to the target outcome, they do not address redundancy and overfitting issues. The maximum relevance and minimum redundancy (MRMR) algorithm aims to reduce the redundancy in datasets, while also identifying the most relevant features and biomarkers to predict the outcome.^31,34^ Furthermore, using the right classification algorithm for the problem is important in order to avoid overfitting by the model.^37^ The classification algorithms that have been used in the past to classify toxicogenomics data include k-nearest neighbor, naïve-Bayes, and support vector machines (SVMs). Support vector machines (SVMs) have been shown to yield reliable and efficient classification performances, while limiting overfitting, particularly for cases where the number of features is higher than the number of samples (as often seen with toxicogenomic data).^29^

Although a few isolated biomarkers have been used for genotoxicity detection in both environmental and human health applications, such as CYP1A1 and CYP1B1,^38^ RAD54,^39^ CYP-R,^38^ A2m, Ca3, Cxcl1, and Cyp8b1,^40^ their correlation with phenotypic genotoxicity endpoints or carcinogenicity has not been quantified. Furthermore, the temporal dependencies of toxicogenomics responses have also not been considered in most cases, since most studies record a snapshot of the responses.^38^ It is still an open research area to identify relevant toxicogenomic-based biomarkers that quantitatively link *in-vitro* responses to regulatory relevant *in-vivo* toxicity endpoints, utilizing the temporal molecular response patterns.

In this study, we applied MRMR feature selection and SVM classification algorithm, to identify an ensemble of biomarkers from temporal toxicogenomic assays, for genotoxicity and carcinogenicity prediction and for bridging to regulatory relevant phenotypic endpoints via AOP. As per the AOP framework, molecular initiating event for DNA damage would link to an adverse outcome of genotoxicity at organism or population level that is relevant to risk assessment.^13,19,41^ We proposed and developed a novel quantitative toxicogenomics assay to evaluate mechanistic genotoxicity through detect and quantifying molecular level changes in proteins involved in known DNA damage repair pathways, to comply with the AOP concept.^2,42-43^ The selected key proteins involved in all known DNA damage and repair stress response pathways are conserved among yeast and other eukaryotes including human, therefore is expected to capture AOP molecular effects at sub-cytotoxic dose levels that lead to phenotypic changes and adverse outcome.^2,19,44^ The protein expression changes, in exposure to each chemical, are monitored by employing a genotoxicity assay using GFP-tagged yeast reporter stains, covering 38 selected protein biomarkers indicative of all the seven known DNA damage repair pathways as discussed above.^2,45^ Two separate case studies — i) *in-vivo* rodent carcinogenicity and ii) Ames test-based genotoxicity prediction — are performed to identify the biomarker-ensembles for chemically-induced genotoxicity and carcinogenicity endpoint prediction. For each case study, 20 chemicals are selected that include model genotoxic compounds with reported endpoints and negative control without any reported genotoxic effects. Both *in-vivo* rodent carcinogenicity and Ames based genotoxicity are among the most widely used endpoints for genotoxicity assessment and they are being used in the National Toxicology Program^46^ and Carcinogenic Potency Database (CPDB)^47^. The performance of the prediction models is evaluated by estimating the area under the receiver operating characteristics curve (AUC), as well as the classification accuracy, sensitivity, and specificity. The relationship between the number and identities of top-ranked biomarker selection and the prediction performances are assessed.

## METHODOLOGY

### Materials

A time-series toxicogenomic assay of 20 chemicals is evaluated in the current study, including model genotoxic compounds and negative control without any reported genotoxic effect. Details of the chemicals are provided in **Table S1**. Two types of genotoxicity endpoints are investigated, including i) *in-vivo* rodent carcinogenicity and ii) Ames genotoxicity assay. Both carcinogenicity and genotoxicity endpoint data is collected from the existing literature and are summarized in **Table S1**.^2,48^

### Quantitative Toxicogenomic Assay for Genotoxicity Assessment

The proteomic based toxicogenomic assay employs a library of 38 in-frame GFP-fused reporter strains (key proteins) of *Saccharomyces cerevisiae* (Invitrogen, no. 95702, ATCC 201388), covering all known recognized DNA damage repair pathways, as reported in our previous work (**Table S2**).^2,49^ The reporter strains are constructed by oligonucleotide-directed homologous recombination to tag each open reading frame (ORF) with Aequrea victoria GFP (S65T) in its chromosomal location at the 3’ end. The assay library measures the expression of full-length, chromosomally tagged green fluorescent protein fusion proteins,^50^ where the GFP signal represents the protein expression, directly. The altered expression level is measured for 2-hr at every 5 minute, which is then integrated over the full exposure period to obtain the quantitative toxicity index — Protein Expression Level Index (PELI). The details of the proteomics assay, when using GFP-tagged yeast cells, were described in the Supporting Information (Text S1) and also in our previous reports.^2,49,51^

### Scoring Criteria to Rank the Biomarkers

The selection of biomarkers is based on their contribution to differentiate protein expression level among the genotoxicity-negative and -positive chemicals. The maximum relevance is used as a scoring measure, which is quantified by estimating t-stat (Text S1) to find the relevance of biomarkers to a target outcome. The second scoring measure, maximum relevance minimum redundancy (MRMR), is applied to reduce the redundancy from the relevant biomarkers and quantified *via* penalizing the relevance for collinearity with other biomarkers in the library.

#### Maximum Relevance

Maximum relevance measures how the biomarkers are relevant to identify the genotoxicity-positive chemicals. t-stat criterion is used as a measure of the relevance.^31^ t-stat measures the difference between the mean of the features in genotoxicity-negative, and genotoxicity-positive class. The higher the t-stat value the higher the differences in average protein expression level between the two classes — genotoxicity-positive and -negative categories. Hence, the objective function of selecting the most significant biomarker is to maximize the relevance, *V*_*t*_.

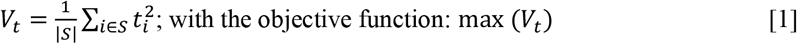

where, *V*_*t*_ measures the relevance of feature *i*,

*t*_*i*_ is the t-stat of the protein *i* (Text S2), and

*S* is the number of features in the dataset.

#### Maximum Relevance Minimum Redundancy

Application of the maximum relevance criteria may identify multiple biomarkers that are correlated with each other. To reduce the redundancy, it is expected that, the selected biomarkers should contain only the uncorrelated proteins since multiple correlated proteins do not provide any additional information regarding the chemical class.^31,52^ Co-linearity can be measured using Pearson correlation between the genes that quantify redundancy, as described in the following.

##### Minimum redundancy

The redundancy (*W*_*c*_) of the biomarkers can be measured by the Pearson correlation of a biomarker with the rest of the biomarkers.

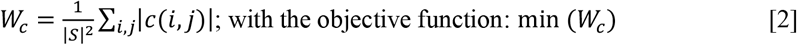

where, *c*(*i, j*) is the correlation between proteins *i* and *j* (Text S2).

By combining the maximum relevance (**Equation 1**) and minimum redundancy (**Equation 2**), in two separate ways, two different maximum relevance minimum redundancy (MRMR) scoring criteria can be computed. They are:

i. MRMR-TCD (t-stat correlation difference criterion): In MRMR-TCD scoring method the relevance and redundancy are combined using difference, where the objective function is: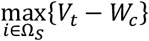. and
ii. MRMR-TCQ (t-stat correlation quotient criterion): MRMR-TCQ score is measured by combining the relevance and redundancy using quotient and the objective function is: 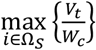. Since MRMR-TCQ apply greater penalty on redundancy than the MRMR-TCD, it has the potential to select the biomarkers that can generate models with better prediction accuracies.^31^

By utilizing the scoring measures, mentioned here, three scoring criteria — t-stat, MRMR-TCD, and MRMR-TCQ — are adopted to find the ranks of the biomarkers. The rank measure of all the scoring methods are calculated using a 10-fold cross-validation (CV). The dataset is randomly divided into 10 equal sized subsets and nine of the ten subsets are used as the training data to get the score. This process is repeated 10 times with different training dataset, achieved by leaving out each of the ten subsets exactly once from the training dataset. The overall score, which is used as the ranking measure, is calculated as the average of the scores from the ten folds.

### Classification Algorithm

The prediction ability of the selected top-ranked biomarkers is evaluated by fitting a classification model, using Support Vector Machine (SVM) with 10-fold CV as the classification algorithm. SVM is chosen due to its power of avoiding overfitting when the feature space is very large.^29^ In a 10-fold CV, the data are randomly divided into 10 equal sized folds, and among them 9 folds are used as the training dataset to generate classification model with the remaining fold as prediction test.^53^ The classification is conducted for 10 times by using each subset as the test dataset exactly once and the remaining nine subsets as training dataset.

#### Classification Performance Measuring Criteria

Receiver operating characteristics (ROC)^54^ curve of the classification models are obtained from the fitted models, and the area under the ROC curve (AUC) is estimated to evaluate the classification model efficiency.^55-56^ The classification accuracy, sensitivity, and specificity are also calculated as additional performance measures. Accuracy is defined as the proportion of all the samples (both true positives and true negatives) that are identified correctly among the total number of samples.^57^ Sensitivity measures the proportion of positives that are correctly identified and specificity measures the proportion of negatives that are correctly identified as such.^58^ The accuracy, sensitivity, and specificity of the models are estimated from the test datasets of all 10 folds, mentioned above, and average of the classification models of the 10 folds are reported as the CV-accuracy, -sensitivity, and -specificity. Both MRMR feature selection and SVM classification algorithm are implemented in MATLAB 2017a (Mathworks, Natick, MA).

### Gene Ontology

Gene ontology (GO) analysis is performed using the Functional Specification resource, FunSpec,^59^ to determine the significantly represented functional categories that are associated with the selected top-ranked biomarkers. The significant biological categories are obtained using a hypergeometric distribution with a p-value threshold of 0.01 — which represent that, the association between a given gene set and a given functional category does not occur at random.^60^

## RESULTS AND DISCUSSION

### Identification of Toxicogenomics-based Biomarkers for *In-vivo* Carcinogenicity Prediction

The most relevant protein biomarkers are identified based on their rank measures and ability to differentiate the altered expression level between the carcinogenicity-positive and -negative compounds. Three separate scores according to the three ranking criteria, namely t-stat, MRMR-TCD and MRMR-TCQ, as described in the methods section, were used. **Figure 1** shows the most relevant biomarkers that have higher scores and assumingly higher relevancy to *in-vivo* carcinogenicity. The higher t-stat score represents a larger difference of the altered expression level between the average biomarker-responses in the carcinogenicity-positive and -negative chemicals (**Figure 1-a**). However, presence of collinearity with other potential biomarkers may result in redundancy among the top-ranked biomarkers, with higher scores. The MRMR scoring criteria helps to eliminate redundancy by penalizing the co-linearity for redundancy elimination. Two MRMR-based ranking criteria (MRMR-TCD, and MRMR-TCQ) rank the proteins after penalizing the t-stat score for the presence of collinearity, which is used as a measure of redundancy (**Figure 1-b, c**). Due to the penalization for redundancy, the rank of a few of the biomarkers are different for the MRMR-based ranking methods than those for the t-stat-based ranking. As the relevance is divided by the redundancy in the MRMR-TCQ, the penalty for collinearity is high in this case. Subsequently, the margin between the high ranked biomarkers and low ranked biomarkers are higher in the case of MRMR-TCQ ranking than the results obtained from MRMR-TCD score. Disregard the ranking algorithms used, the top-ranked five protein biomarkers are identified to be the same for all three scoring methods, although their rank sequences varied. These five proteins, namely NTG2, RAD34, RAD27, MSH2, and YKU70, can be considered to have high relevance with very little redundancy, which make them suitable as potential biomarkers for *in-vivo* carcinogenicity prediction.

**Figure 1.**
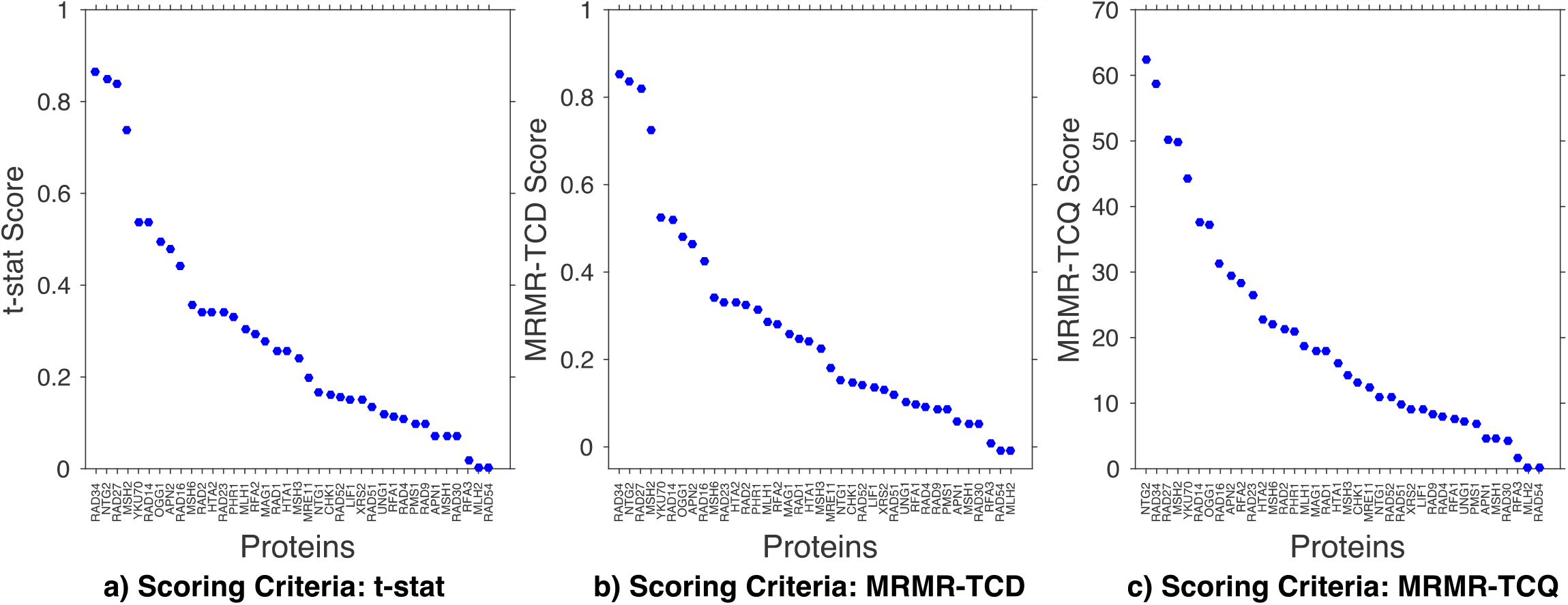
Scores of the biomarker based on different scoring criteria for Carcinogenicity endpoint prediction. The biomarkers are sorted in descending order of their score. The higher the score, the more important the biomarker is. a) Ranking criterion: t-stat. This ranking method is based on maximum relevance alone. The t-stat measures the significant differences between the mean of the biomarker feature (in this specific case: PELI) in the positive and negative carcinogenic chemicals. The higher the difference in mean of the biomarkers in the two classes, the more important the biomarker is. b) Ranking criterion: MRMR-TCD. This ranking criterion is based on the maximum relevance and minimum redundancy. The t-stat measures the relevance to the endpoint. Collinearity among different biomarkers can measure the redundancy of the biomarker. In the MRMR-TCD ranking method, average correlation of the biomarker is subtracted from the t-stat score to remove the redundancy. c) Ranking criterion: MRMR-TCQ. This criterion is also based on the maximum relevance and minimum redundancy. The t-stat is divided by the average correlation, to get the maximum relevance and minimum score.

A heat-map of the molecular genotoxicity quantifier, PELI (protein effect level index) values, shows the differential expression pattern of the top-ranked biomarkers among carcinogenicity-positive and -negative chemicals (**Figure 2**). The preferred top-ranked biomarker(s) exhibited significantly higher altered protein expression level in carcinogenicity-positive chemicals, whereas significantly less or negligible (below detection limit PELI < 1.5)^2,49^ altered expression level for the carcinogenicity-negative chemicals. In contrast, the bottom-ranked protein had similar average PELI values in both carcinogenicity-positive and -negative chemicals.

**Figure 2.**
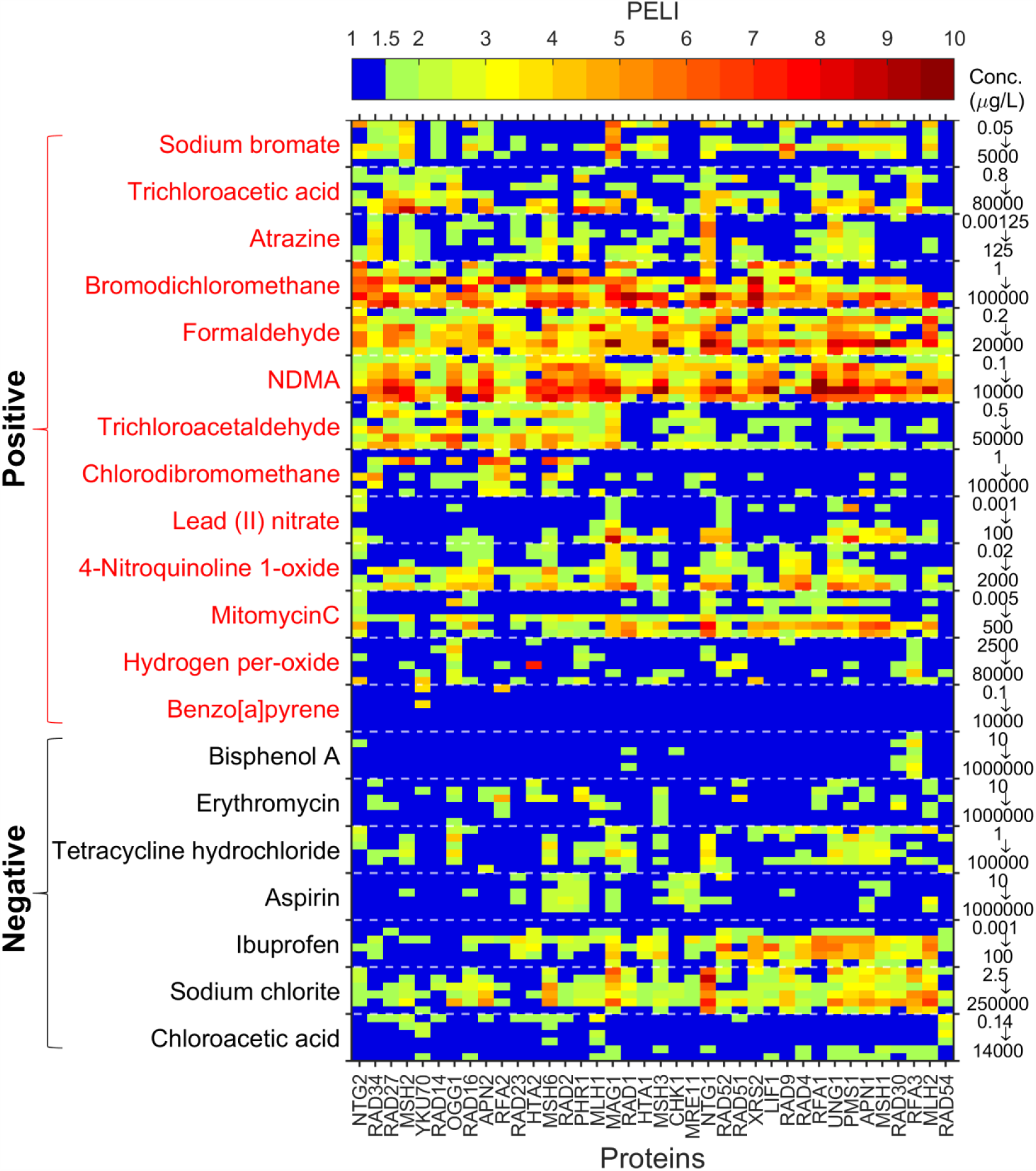
Heatmap of the PELI values (colormap as indicated by the color bar) of all the protein biomarkers covering all known DNA damage and repair pathways in yeast (*Saccharomyces cerevisiae*), upon exposure to 20 chemicals (rows). For each chemical, responses to 6 doses are presented with concentrations vary from lowest to highest from top to bottom (noted on the right-side of vertical axis, all concentrations are listed in **Table S1**). The Biomarkers (columns) are sorted (from left to right) as per their ranks based on the MRMR-TCQ score (**Figure 1-c**). The carcinogenicity-positive chemicals (rows) are labeled in red color, and the carcinogenicity-negative chemicals are labeled in black color.

#### Performance evaluation of the top-ranked biomarkers for in-vivo *carcinogenicity prediction*

We evaluate the number and the ability of the top-ranked protein biomarkers as biomarker-ensemble to predict chemical-induced *in-vivo* carcinogenicity. Support vector machine (SVM) is used to classify and predict the carcinogenetic compounds with varying number of top-ranked biomarkers. The area under the receiver operating characteristics curve (AUC) is used as the classifier performance evaluation criterion, which indicates the stability of classification model with AUC value closer to 1 representing a stable classifier.^61^ For the *in-vivo* carcinogenicity prediction, AUC is 0.81 with SVM model using the top-five ranked biomarkers (**Table 1**). The increase in the number of biomarkers in the classifier model, as expected, leads to increase in the AUC. The maximum AUC value of the models is 0.88, when all the 38 proteins in the library are used.

**Table 1.** Prediction model performance of the two case-studies. The model performance measures include AUC, classification accuracy, sensitivity, and specificity. The prediction models are generated with top-ranked biomarkers as well as with all the proteins in the assay library.

| Number of Biomarkers utilized in the classification model | AUC | Classification Performance Metric |  |  |
| --- | --- | --- | --- | --- |
|  |  | Accuracy | Sensitivity | Specificity |
| *In-vivo* carcinogenicity prediction models |  |  |  |  |
| Top Five | 0.81 | 76% | 74% | 79% |
| All | 0.88 | 83% | 81% | 86% |
| Ames genotoxicity prediction models |  |  |  |  |
| Top Five | 0.75 | 70% | 63% | 81% |
| All | 0.84 | 78% | 79% | 75% |

Furthermore, the cross-validated accuracy, sensitivity, and specificity of the 10-fold models are estimated as additional performance measuring criteria. Top-ranked five biomarkers can obtain 76% accuracy for the carcinogenicity prediction, while the classification model with all the proteins can achieve 83% accuracy. The sensitivity and specificities also increase with the increase of the number of biomarkers in the classifier models. For example, sensitivity of the classification model is 74% for the prediction model with top five biomarkers and it reached 81% with all the proteins. Specificity increases from 79%, for the model with the top five biomarkers, to 86% for the model where all the proteins are used as features. Relatively high CV-accuracies, which are measured from the prediction of the test dataset, indicate the robustness of the prediction model obtained through the SVM classification model. Although, as expected, higher number of features in the classification models improves the accuracy and the stability, the top five identified most relevant biomarkers can achieve classification accuracy of ∼75%, implying a possible cost- and time-effective *in-vivo* carcinogenicity screening tool with the time-series toxicogenomic assay.

### Identification of Toxicogenomics-based Biomarkers for Ames Genotoxicity Prediction

We performed a second case study, where we attempted to identify the protein biomarkers for the Ames genotoxicity endpoint prediction. The most important biomarkers are identified from their ranking scores, which are based on their difference in altered protein expression level between the Ames genotoxicity-positive and -negative compounds. The three separate scores of each biomarker, obtained from the three ranking criteria (t-stat, MRMR-TCD and MRMR-TCQ), are presented in **Figure 3**. The higher difference between the altered expression level of a protein in the genotoxicity-positive and -negative chemicals results in relatively higher t-stat scores (**Figure 3-a**). Hence, the top-ranked proteins, with higher t-stat scores, have the higher relevance to differentiate Ames genotoxicity-positive chemicals from the genotoxicity-negative chemicals. The MRMR-TCD and MRMR-TCQ based scores differ from the t-stat score, especially for the proteins that possess collinearity with other proteins in the assay (**Figure 3-b, c**). However, the identified top-ranked five protein biomarkers remain the same irrespective of the ranking methods employed, although their rank sequences varied. These five proteins, namely APN2, RFA2, NTG2, RAD2, and MSH6, are considered as potential biomarkers for the Ames genotoxicity prediction.

**Figure 3.**
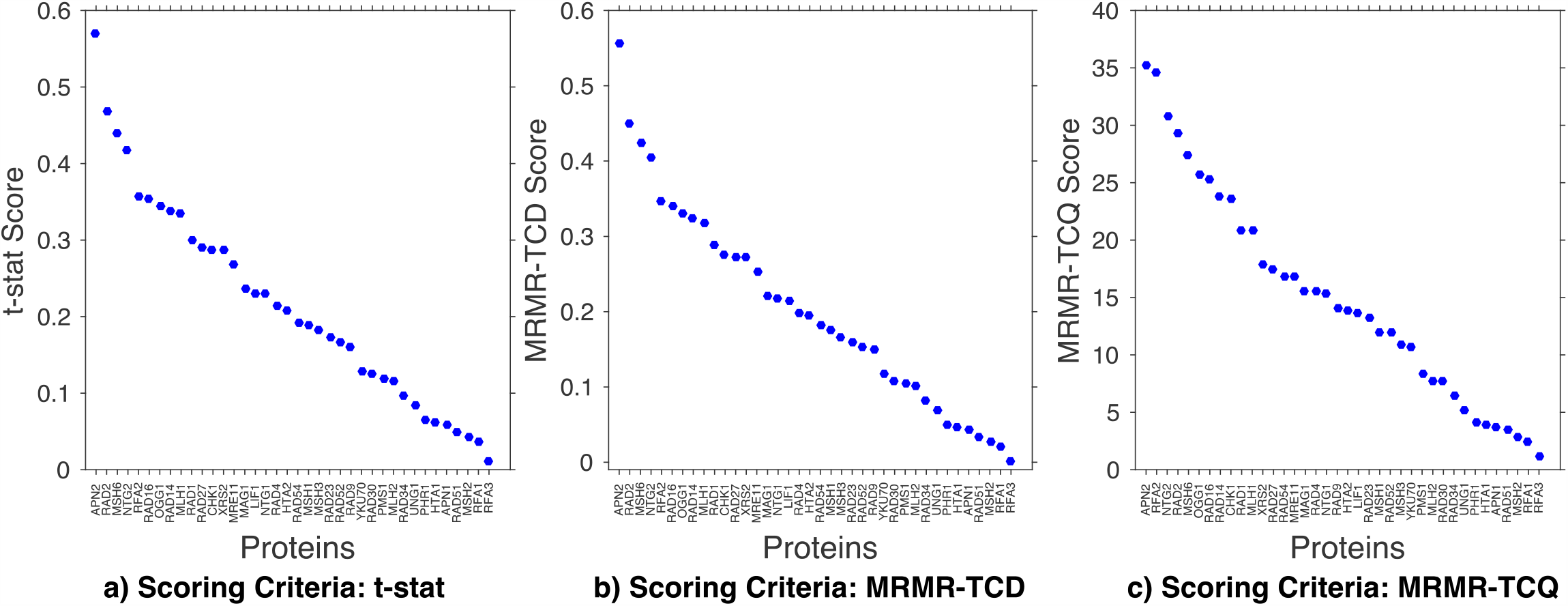
Scores of the biomarker based on different scoring criteria for Ames-genotoxicity assay prediction. The biomarkers are sorted in descending order of their score. The higher the score, the more important the biomarker is. **a)** Ranking criterion: t-stat. This ranking method is based on the maximum relevance. The t-stat measures the significant differences between the mean of the biomarker feature (in this specific case: PELI) in the chemicals with Ames assay positive and negative responses. The higher the difference in mean of the biomarkers in the two classes, the more important the biomarker is. **b)** Ranking criterion: MRMR-TCD. This ranking criterion is based on the maximum relevance and minimum redundancy. The t-stat measures the relevance to the endpoint. Collinearity among different biomarkers can measure the redundancy of the biomarker. In the MRMR-TCD ranking method, average correlation of the biomarker is subtracted from the t-stat score to remove the redundancy. **c)** Ranking criterion: MRMR-TCQ. This criterion is also based on the maximum relevance and minimum redundancy. The t-stat is divided by the average correlation, to get the maximum relevance and minimum score.

A heat-map of the molecular genotoxicity quantifier, PELI, further shows the distinct expression profiles of the top ranked biomarkers in the genotoxicity-positive and -negative chemicals (**Figure 4**). The top-ranked biomarkers show relatively higher protein expression level in genotoxicity-positive chemicals than the expression level in genotoxicity-negative chemicals that are often below toxicity threshold (PELI < 1.5).^2,49^ For the bottom-ranked proteins, the difference in expression levels across genotoxicity-positive and -negative chemicals decreases, which further suggests their inadequacy to separate genotoxicity-positive chemicals from the genotoxicity-negative chemicals.

**Figure 4.**
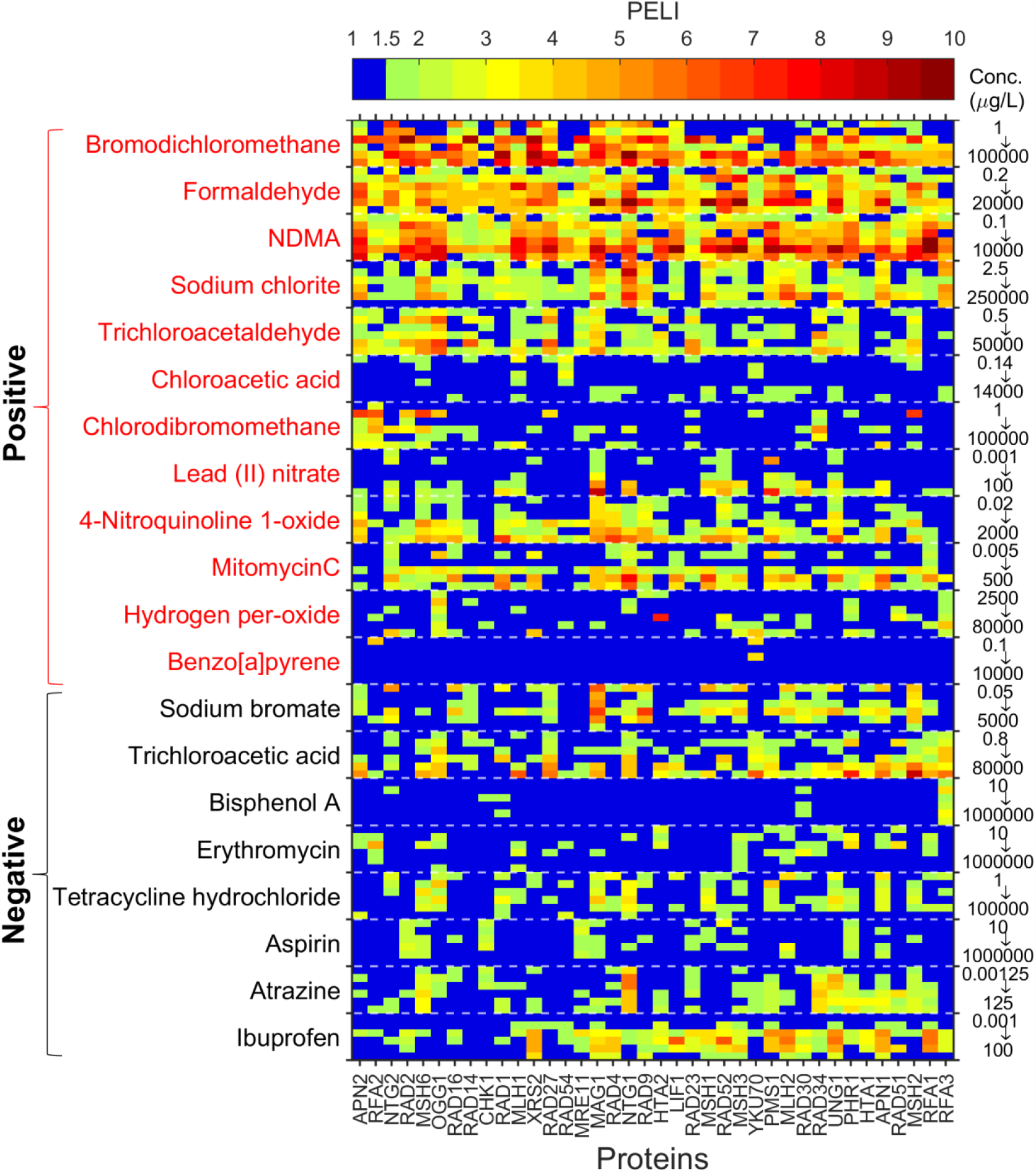
Heatmap of the PELI values (colormap as indicated by the color bar) of all the protein biomarkers covering all known DNA damage and repair pathways in yeast (*Saccharomyces cerevisiae*), upon exposure to 20 chemicals (rows). For each chemical, responses to 6 doses are presented with concentrations vary from lowest to highest from top to bottom (noted on the right-side of vertical axis, all concentrations are listed in **Table S1**).The Biomarkers (columns) are sorted (from left to right) as per their ranks based on the MRMR-TCQ score (**Figure 3-c**). The Ames genotoxicity-positive chemicals (rows) are labeled in red color, and the genotoxicity-negative chemicals are labeled in black color.

#### Performance evaluation of the top-ranked biomarkers for Ames genotoxicity prediction

The prediction ability of the top-ranked biomarker-ensemble is evaluated by using the SVM to classify and predict compounds with Ames genotoxicity with varying number of top-ranked biomarkers. The classifier performances are evaluated by estimating the AUC of the SVM models (**Table 1**). AUC of the model with top-ranked five biomarkers is 0.75 and that of the model with all 38 proteins is 0.84. AUC values of both the models represent stable classification models, since the AUC value equal to 0.5 indicates a random classification model and closer to 1 represents a stable model^61^. As expected, the prediction model with all the proteins has higher AUC than the model with the top-five biomarkers.

The estimated model accuracy, sensitivity, and specificity provide additional measures of the prediction performance. Using only the top-ranked five biomarkers can achieve 70% accuracy, which increased to 78% for the model with all 38 proteins. The sensitivity (63% for top-five biomarkers and 79% for all 38 proteins) of the prediction model also increases with the increase in number of biomarkers in the classification model. Though the specificity of the prediction model with top-ranked five biomarkers (81%) is higher than that of the model with all 38 proteins (75%), the specificity of the model with all the protein biomarkers are closer to its accuracy representing relatively low model-bias. It can be inferred that, though higher number of biomarkers results in improvement of prediction, the top-ranked five biomarkers can achieve relatively high prediction accuracy (70%). Therefore, possible application of time-series toxicogenomic assay with the most relevant top-ranked biomarkers may provide a balance of cost- and time-effective Ames genotoxicity screening for Ames effect assessment and for further in-depth studies.

### Gene Ontology Analysis Reveals the Biological Processes and Molecular Functions of the Top-Ranked Biomarkers

Gene ontology (GO)^60^ analysis is conducted to reveal the biological processes and molecular functions of the top-ranked biomarkers. Biological processes and molecular functions that are significantly associated with the top-ranked five biomarkers are determined using the hypergeometric distribution with a threshold p-value of 0.01, which indicates that the associations between the biomarkers and the GO terms are not at random.^59-60^

#### Biological processes and molecular functions of the biomarkers for carcinogenicity prediction

GO analysis reveals that, the most significantly enriched biological processes of the top-ranked five biomarkers (RAD34, RAD27, YKU70, NTG2, MSH2), for the *in-vivo* carcinogenicity prediction, are associated with DNA repair and response to DNA damage stimulus (**Table S3**). Additionally, none of the biomarkers is annotated with the functional categories that represent other types of response pathways. RAD27 and YKU70 are associated with double strand break repair. DNA recombination involves YKU70 and MSH2. The biomarkers are also relevant to the base-excision repair and double-strand break.

The significantly enriched molecular functions associated with the top-ranked five biomarkers are also relevant to the DNA damage and repair activities. Damaged DNA binding is associated with RAD34 and YKU70. Endonuclease activity is associated with RAD27, and NTG2. The other relevant molecular functions, associated with at least one of the top-ranked five biomarkers, include single base insertion or deletion binding, DNA binding, DNA insertion of deletion binding, double-strand/single-strand DNA junction binding, and mismatched DNA binding (**Table S3**). The top-ranked five biomarkers for the *in-vivo* carcinogenicity prediction mainly focused on double strand break repair and DNA recombination, which result in a severe type of DNA damage and may lead to genomic instability and cancer.^62-63^ This suggests that the selected biomarker-ensemble have the potential to identify severe DNA damage and relevant carcinogenic potency of a chemical.

#### Biological processes and molecular functions of the biomarkers for Ames genotoxicity prediction

The most significantly enriched biological processes of the top-ranked five biomarkers for the Ames genotoxicity prediction are also revealed *via* GO analysis. These biological functional categories are also associated with DNA damage and repair activities, and do not show any association with other stress response pathways. The top-ranked five biomarkers (APN2, RFA2, NTG2, RAD2, MSH6) are associated with the DNA repair process and four of them (APN2, NTG2, RAD2, MSH6) are related to the response to DNA damage stimulus biological process. Other biological processes, which show significant association with at least one of the top-ranked biomarkers, include base-excision repair, nucleotide-excision repair, DNA unwinding involved in replication, and double-strand break repair via homologous recombination (**Table S3**).

The most significantly enriched molecular functions of the top-ranked five biomarkers are also related to DNA damage and repair activities. Endonuclease activity is associated with three of the top-five biomarkers (APN2, RAD2, NTG2). DNA binding molecular function is also associated with three biomarkers, which are APN2, MSH6, and RAD2. Other relevant molecular functions that are significantly associated with at least one of the top-five biomarkers include DNA (apurinic or apyrimidinic site) lyase activity, single-stranded DNA binding, double-stranded DNA specific 3’-5’ exodeoxyribonuclease activity, single base insertion or deletion binding, four-way junction DNA binding, DNA binding, and mismatched DNA binding. The selected top-ranked biomarkers for Ames genotoxicity prediction are associated with base- and nucleotide-excision repair. Since Ames genotoxicity test only detects the frame-shift mutation and base-pair substitution,^64-65^ it captures certain genotoxicity effects, and results in the selection of different biomarker-ensemble than those in the *in-vivo* carcinogenicity prediction case-study.

### Environmental Implications

Presence of numerous contaminants from various (e.g., wastewater effluent, industrial wastewater discharge, oil and chemical spills, urban runoff, agricultural runoff) sources make the risk assessment and monitoring challenging.^17^ High-throughput in vitro assays and AOP-based approach is promising for the assessment of health and ecotoxicological risks from exposure to pollutants and their mixtures.^17-21^ Establishment of a quantitative AOP framework through integration of computational modeling with *in-vitro* assays would require identification and selection of relevant biomarkers from appropriate bioassays that link to risk assessment pathways and phenotypic impacts.^18,21,66^ Current toxicomics approach still mostly rely on large number of redundant markers without pre-selection or ranking;^3,51,67^ therefore, selection of relevant biomarkers with minimal redundancy would reduce the number of markers to be monitored and reduce the cost, time, and complexity of the toxicity risk monitoring. The method developed in this study will help to fill in the knowledge gap in phenotypic anchoring and predictive toxicology, and contribute to the progress in the implementation of tox 21 vision for environmental and health applications.

## SUPPORTING INFORMATION

Tables listing the selected concentration ranges and sources of the chemicals, *in-vivo* carcinogenicity and Ames genotoxicity endpoints along with their sources, description and data processing steps of the yeast proteomic library used in this study, and the summary of gene ontology analysis of the selected biomarkers are provided in the Supporting Information (SI). This material is available free of charge via the Internet at http://pubs.acs.org.

## ACKNOWLEDGEMENTS

This study is funded by National Science Foundation (NSF, CBET-1437257, CBET-1810769, and IIS-1546428), National Institute of Environmental Health Sciences (NIEHS, PROTECT 3P42ES017109 and CRECE 5P50ES026049), and Environmental Protection Agency (EPA, CRECE 83615501).

