## supporting information for "Machine Learning-based Biomarkers Identification and Validation from Toxicogenomics - Bridging to Regulatory Relevant Phenotypic Endpoints"

No. of Pages: 12

No. of Tables: 3

No. of Text Sections: 2

**Table of Content**

**Table S2. DNA damage type, corresponding repair pathways and biomarkers selected for the molecular genotoxicity assay using GFP-tagged yeast cells.**

[**Table S3. Significantly represented biological categories based on the GO analysis of the top‑ranked five biomarkers that are selected for phenotypic toxicity prediction. The significant GO‑term are identified using the hypergeometric probability distribution with p‑value < 0.01.**](#_Toc5316153)

Text S1. Quantitative Toxicogenomic Assay for Genotoxicity Assessment

Text S2. Equations for the calculation of t-stat and Pearson correlation

Table S1. List of chemicals, concentration range, the *in-vivo* carcinogenicity and Ames genotoxicity endpoints of the chemicals selected in the study.

| **Chemical Name** | **Source**  **(cas#; cat#)** | **Concentration range ^‡^**  **6 doses with 10x dilution**  **(mg/L)** | ***In-vivo* Carcinogenicity ^$^** | **Ames genotoxicity ^$^** |
| --- | --- | --- | --- | --- |
| 4-Nitroquinoline 1-oxide | Acros Organics  (56-57-5; AC20379‑2500) | 0.00002 - 2 | Positive ^e^ | Positive ^n^ |
| Aspirin | Fisher Scientific  (50-78-2; S79881) | 0.01 - 1000 | Negative ^d^ | Negative ^k^ |
| Atrazine | AccuStandard  (1912-24-9; P-005S) | 0.00000125 - 0.125 | Positive ^d^ | Negative ^l^ |
| Benzo[a]pyrene | Sigma-Aldrich  (50-32-8; B1760‑250mg) | 0.0001 - 10 | Positive ^d^ | Positive ^m^ |
| Bisphenol A | Sigma-Aldrich  (80-05-7; 239658-50G) | 0.01 - 1000 | Negative ^d^ | Negative ^p^ |
| Bromodichloromethane | ACROS Organics  (75-27-4; 160700100) | 0.001 - 100 | Positive ^c^ | Positive ^c^ |
| Chloroacetic acid | ACROS Organics  (79-11-8; 108512500) | 0.00014 - 14 | Negative ^c^ | Positive ^c^ |
| Chlorodibromomethane | ALFA AESAR  (124-48-1; A16938-06) | 0.001 - 100 | Positive ^c^ | Positive ^c^ |
| Erythromycin | MP Biomedicals  (190197; 114-07-8) | 0.01 - 1000 | Negative ^d^ | Negative ^q^ |
| Formaldehyde | 37% by Weight, Fisher Chemical  (50-00-0; F79-500) | 0.0002 - 20 | Positive ^c^ | Positive ^h^ |
| Hydrogen per-oxide | Sigma-Aldrich  (7722-84-1; 349887‑500ml) | 2.5 – 80  (2x dilution) | Positive ^r^ | Positive ^n^ |
| Ibuprofen | Acros Organics  (15687-27-1;  AC258610100) | 0.000001 - 0.1 | Negative ^d^ | Negative ^j^ |
| Lead (II) nitrate | Acros Organics  (10099-74-8;  AC196071000) | 0.000001 - 0.1 | Positive ^s^ | Positive ^t^ |
| MitomycinC | RPI corp.  (50-07-7; M92010-0.002) | 0.000005 - 0.5 | Positive ^d^ | Positive ^n^ |
| NDMA  (n-nitrosodimethylamine) | CHEM SERVICE INC  (62-75-9; N‑12572‑250MG) | 0.0001 - 10 | Positive ^c^ | Positive ^g^ |
| Sodium bromate | EM Science  (7789-38-0; SX0385-1) | 0.00005 - 5 | Positive ^c^ | Negative ^h^ |
| Sodium chlorite | ACROS Organics  (7758-19-2;223235000) | 0.0025 - 250 | Negative ^c^ | Positive ^i^ |
| Tetracycline hydrochloride | Genlantis Inc  (64-75-5; M160025) | 0.001 - 100 | Negative ^d^ | Negative ^u^ |
| Trichloroacetaldehyde | Waterstone Pharmaceuticals, Inc.  (75-87-6; WS51847-1G) | 0.0005 - 50 | Positive ^c^ | Positive ^c^ |
| Trichloroacetic acid | Fisher chemical  (76-03-9; A322-100) | 0.0008 - 80 | Positive ^c^ | Negative ^c^ |

**^‡^** The highest concentration for each chemical is sub-cytotoxic concentration with over 95% cell survival rate from a 24 h exposure (LC5) experiment. See details in previous studies.^1-2^

^$^ The *in-vivo* carcinogenicity and the Ames genotoxicity endpoint data are collected from the literature. The data sources are:

^c^ Richardson *et al.* (2007)^3^, ^d^ Gold *et al.* (2000)^4^, ^e^ Shirasu & Ohta (1963)^5^, ^g^ Stalter *et al.* (2016)^6^, ^h^ HSDB (2018)^7^, ^i^ Kurokawa *et al.* (1986)^8^, ^j^ Oldham *et al.* (1986)^9^, ^k^ Jasiewicz & Richardson (1987)^10^, ^l^ Della Croce *et al.* (1996)^11^, ^m^ Phillipson & Ioannides (1989)^12^ , ^n^ Nakamura *et al.* (1987)^13^, ^p^ Fic *et al.* (2013)^14^, ^q^ Isidori *et al.* (2005)^15^, ^r^ Reifferscheid & Heil (1996)^16^, ^s^ García‑Lestón *et al.* (2010)^17^, ^t^ Rank & Nielsen (1994)^18^, and ^u^ NTP (1982)^19^.

Table S2. DNA damage type, corresponding repair pathways and biomarkers selected for the molecular genotoxicity assay using GFP-tagged yeast cells.^1-2,20-21^

| **DNA damage** | **Repair pathway** | | **Proteins selected in the assay** |
| --- | --- | --- | --- |
| general damage | DNA damage signaling (DDS) | | CHK1, RAD9 |
| DNA lesions | translesion synthesis (TLS) | | RAD30 |
| base alkylation | direct reversal repair (DRR) | | PHR1 |
| base oxidation  base alkylation and deamination  single strand break | base excision repair (BER) | | OGG1  NTG1, NTG2, UNG1, MAG1, RAD27, APN1, APN2 |
| cross links  pyrimidine dimers  bulky adduct | nucleotide excision repair (NER) | | RAD1, RAD2, RAD4, RAD14, RAD16, RAD23, RAD34 |
| mismatches | mismatch repair (MMR) | | MSH1, MSH2, MSH3, MSH6, PMS1, MLH1, MLH2 |
| double strand break (DSB) | DSB repair | general response to DSB | XRS2, MRE11 |
|  |  | homologous recombination (HR) | RFA1, RFA2, RFA3, RAD51, RAD52, RAD54, HTA1, HTA2 |
|  |  | non-homologous end joining (NHEJ) | LIF1, YKU70 |

Table S3. Significantly represented biological categories based on the GO analysis of the top‑ranked five biomarkers that are selected for phenotypic toxicity prediction. The significant GO‑term are identified using the hypergeometric probability distribution with p‑value < 0.01.

| **Categories** | **GO ID** | **Biomarkers** |
| --- | --- | --- |
| **Biomarkers for *in-vivo*** **carcinogenicity prediction** | | |
| *Biological Process* | | |
| DNA repair | 0006281 | RAD34, RAD27, YKU70, NTG2, MSH2 |
| Response to DNA damage stimulus | 0006974 | RAD34, RAD27, YKU70, NTG2, MSH2 |
| Double-strand break repair via nonhomologous end joining | 0006303 | RAD27, YKU70, |
| DNA recombination | 0006310 | YKU70, MSH2 |
| Base-excision repair, base-free sugar-phosphate removal | 0006286 | RAD27 |
| Base-excision repair, AP site formation | 0006285 | NTG2 |
| Double-strand break repair via homologous recombination | 0000724 | RAD27 |
| *Molecular Function* | | |
| Damaged DNA binding | 0003684 | RAD34, YKU70 |
| Endonuclease activity | 0004519 | RAD27, NTG2 |
| Single base insertion or deletion binding | 0032138 | MSH2 |
| Double-strand/single-strand DNA junction binding | 0000406 | MSH2 |
| Oxidized pyrimidine base lesion DNA N-glycosylase activity | 0000703 | NTG2 |
| Y-form DNA binding | 0000403 | MSH2 |
| Guanine/thymine mispair binding | 0032137 | MSH2 |
| DNA binding | 0003677 | RAD27, YKU70, MSH2 |
| DNA insertion or deletion binding | 0032135 | MSH2 |
| 5'-3' exonuclease activity | 0008409 | RAD27 |
| Loop DNA binding | 0000404 | MSH2 |
| Mismatched DNA binding | 0030983 | MSH2 |
| **Biomarkers for Ames genotoxicity prediction** | | |
| *Biological Process* | | |
| DNA repair | 0006281 | APN2, RFA2, NTG2, RAD2, MSH6 |
| Response to DNA damage stimulus | 0006974 | APN2, NTG2, RAD2, MSH6 |
| Base-excision repair | 0006284 | APN2, NTG2 |
| Nucleotide-excision repair | 0006289 | RFA2, RAD2 |
| Nucleotide-excision repair, DNA incision, 3'-to lesion | 0006295 | RAD2 |
| Base-excision repair, AP site formation | 0006285 | NTG2 |
| DNA unwinding involved in replication | 0006268 | RFA2 |
| Double-strand break repair via homologous recombination | 0000724 | RFA2 |
| *Molecular Function* | | |
| Endonuclease activity | 0004519 | APN2, RAD2, NTG2 |
| DNA (apurinic or apyrimidinic site) lyase activity | 0003906 | APN2, NTG2 |
| Single-stranded DNA binding | 0003697 | RAD2, RFA2 |
| Nuclease activity | 0004518 | APN2, RAD2 |
| Oxidized pyrimidine base lesion DNA N-glycosylase activity | 0000703 | NTG2 |
| Double-stranded DNA specific 3'-5' exodeoxyribonuclease activity | 0008311 | APN2 |
| Single base insertion or deletion binding | 0032138 | MSH6 |
| Guanine/thymine mispair binding | 0032137 | MSH6 |
| DNA binding | 0003677 | APN2, MSH6, RAD2 |
| Four-way junction DNA binding | 0000400 | MSH6 |
| Single-stranded DNA specific endodeoxyribonuclease activity | 0000014 | RAD2 |
| Mismatched DNA binding | 0030983 | MSH6 |

Text S1. Quantitative Toxicogenomic Assay for Genotoxicity Assessment

The proteomic based toxicogenomic assay employs a library of 38 in-frame GFP‑fused reporter strains (key proteins) of *Saccharomyces cerevisiae* (Invitrogen, no. 95702, ATCC 201388), covering all known recognized DNA damage repair pathways, as reported in our previous work (**Table S2**).^1,20^ The reporter strains are constructed by oligonucleotide-directed homologous recombination to tag each open reading frame (ORF) with Aequrea victoria GFP (S65T) in its chromosomal location at the 3’ end. The assay library measures the expression of full-length, chromosomally tagged green fluorescent protein fusion proteins,^22^ where the GFP signal represents the protein expression, directly. The altered expression level is measured for 2-hr at every 5 minute. The details of the proteomics assay, when using GFP-tagged yeast cells, were described in our previous reports.^1,20,23^ Briefly, selected yeast strains, cultured in a YPD medium, are grown in clear bottom black 384-well plates in SD medium until they reach the early exponential growth (OD600 of 0.2‑0.4). The chemicals pre-dissolved in PBS, along with blank control (SD medium + 0.25% YPD medium w or w/o chemical), and internal control (SD medium + 0.25% YPD medium + PGK1 strain w or w/o chemical), are added to the well at the selected concentrations. All the chemicals are evaluated across six-log concentration range (except H_2_O_2_, which is evaluated at 2x dilution; concentration ranges of all the chemicals are listed in **Table S1**) with the maximum concentration being sub-cytotoxic (corresponds to >95% survival rate tested by growth inhibition in yeast for 24 hr). Housekeeping gene PGK1 is used as an internal control for plate normalization.^24^ After adding the chemicals and controls, the plates are then placed into a micro plate reader (Synergy H1 Multi-Mode, Biotech, Winooski, VT) for absorbance (OD600 for cell growth) and GFP signal (filters with 485 nm excitation and 535 nm emission for protein expression) measurements for 2-hr at every 5 minute after double orbital shaking (425 cpm) for 1 minute. All tests were performed in dark in triplicate.

*Data Pre-Processing*.

The OD and GFP values of all the wells (w or w/o chemical) are first corrected for any interference due to the media and chemical by subtracting the OD and GFP of corresponding blank controls, with or without chemical. The corrected OD and GFP signals are referred to as *OD_corrected_* and *GFP_corrected_*. The corrected GFP signal (*GFP_corrected_*) is then normalized to the cell numbers in the well (*OD_corrected_*) as *P = GFP_corrected_/OD_corrected_*. The P values are normalized and scaled against the internal control (housekeeping protein PGK1), which is denoted as *Q*. The altered protein expression, relative to untreated control (without chemical), for a given protein ORF *i* in treatment *x* at time point *t* due to chemical exposure, also referred as induction factor *I*, is calculated as:

$I_{i,x,t}=\frac{Q_{i,x,t}}{Q_{i,untreated,t}}$	[1]

where,	*Q_i,x,t_* is the *P* value of ORF *i* at time *t* in treatment *x* (with chemical exposure), normalized against the internal control, and

*Q_i,control,t_* is the *P* value of ORF *i* at time *t* in the untreated control condition (without chemical exposure), normalized against the internal control.

The induction factor, *I* < 1 indicates downregulation and the *I* > 1 indicates upregulation of the biomarkers due to the chemical exposure. The Protein Expression Level Index, PELI, is calculated to aggregate the time series altered expression into a quantitative toxicity index. The chemical induced PELI of the protein is calculated as,^1,20-21^

${PELI}_{ORF,i}= \frac{\int_{t=0}^{t} Idt}{T};where I=1, if I\leq1, else I=I.$	[2]

where, *t* is the exposure time in every time-step, and

*T* is the total exposure time.

Text S2. Equations for the calculation of t-stat and Pearson correlation

*t-stat:*

$t_{i}= \frac{(\bar{g}_{+}-\bar{g}_{-})}{\sqrt{\frac{s_{+}^{2}}{n_{+}} + \frac{s_{-}^{2}}{n_{-}}}}$	[3]

where,	*t_i_* is the t-stat of the protein *i*,

$\bar{g}_{+}$ is the mean of expression of samples in the positive class,

$\bar{g}_{-}$is the mean of the expression of samples in the negative class,

$s_{+}$ is the standard deviation of the positive class,

$s_{-}$ is the standard deviation of negative class,

$n_{+}$ is the number of positive class samples, and

$n_{-}$ is the number of negative class samples.

*Pearson Correlation:*

$c(i,j)=\frac{\sum_{k=1}^{N} (x_{i,k}-\bar{x}_{i})(x_{j,k}-\bar{x}_{j})}{\sqrt{\sum_{k=1}^{N} {(x_{i,k}-\bar{x}_{i})}^{2}}\sqrt{\sum_{k=1}^{N} {(x_{j,k}-\bar{x}_{j})}^{2}}}$	[4]

where,	$c(i,j)$ is the correlation between proteins *i* and *j*

*x_i,k_* is the expression of protein *i* for sample *k*.

*x_j,k_* is the expression of protein *j* for sample *k*,

$\bar{x}_{i}$ is the mean of the expression of protein *i* across all the samples,

$\bar{x}_{j}$ is the mean of the expression of protein *j* across all the samples,

*N* is the number of samples,
